# Molecular and epistatic interactions between pioneer transcription factors shape nucleosome dynamics and cell differentiation

**DOI:** 10.1101/2024.05.27.596047

**Authors:** Rémi-Xavier Coux, Agnès Dubois, Almira Chervova, Nicola Festuccia, Inma Gonzalez, Sandrine Vandormael-Pournin, Michel Cohen-Tannoudji, Pablo Navarro

**Affiliations:** Department of Developmental and Stem Cell Biology, Institut Pasteur, Université Paris Cité, CNRS UMR3738, Paris, France; Epigenomics, Proliferation, and the Identity of Cells Unit; Early Mammalian Development and Stem Cell Biology Group

## Abstract

Pioneer transcription factors (TF) bind nucleosome-embedded DNA motifs to activate new regulatory elements and promote differentiation. However, the complexity, binding dependencies and temporal effects of their action remain unclear. Here, we dissect how the pioneer TF GATA6 triggers Primitive Endoderm (PrE) differentiation from pluripotent cells. We show that transient GATA6 binding exploits accessible regions to decommission active enhancers and promote pluripotency gene silencing. Simultaneously, GATA6 targets closed chromatin and initiates an extensive remodeling culminating in the establishment of fragile nucleosomes flanked by ordered nucleosome arrays and increased accessibility. This is directly enhanced by rapidly expressed PrE TFs (SOX17) and by pluripotency TFs repurposed for differentiation (OCT4/SOX2). Furthermore, GATA6 mediates the replacement of essential nuclear receptors for PrE differentiation, from ESRRB to ESRRA. Therefore, pioneer TFs orchestrate a complex gene regulatory network involving many if not all available pioneer TFs, including those required to support the original identity of differentiating cells.

## Introduction

The acquisition of new cell identities during differentiation processes requires the activation of genes and their regulatory elements^**1**^ even when they are tightly packed into chromatin^**2**^ and inaccessible to many Transcription Factors (TFs)^**3**^. Indeed, the nucleosome, the basic structural unit of the chromatin constituted of around 146 bases of DNA wrapped around an octamer of histone proteins^**4**^, represents a barrier for most TFs to bind^**5**^. Important exceptions exist, however, referred to as pioneer TFs that bind to their cognate motifs even when packed into nucleosomes^**6**^. By licensing otherwise closed and inactive regulatory elements, pioneer TFs initiate large reprogramming events associated with the establishment of the transcription profiles driving and assisting cell differentiation^**7-9**^.

Members of the GATA family have long been shown to display properties of pioneer TFs^**10**^. GATA1 was first shown to bind *in vitro* to a nucleosome harboring binding motifs, leading to the destabilization of histone-DNA contacts and to increased sensitivity to digestion by nucleases^**11**^. Later, in the seminal work coining the term pioneer TF, another member with a recognized role in embryonic^**10,12**^ and extra-embryonic endoderm differentiation^**13,14**^, GATA4, was shown to bind and open nucleosomal arrays *in vitro*^**15**^. Subsequently, other members, including GATA6, were shown to bind nucleosome arrays *in vitro* or suggested to do so in cellular contexts where they trigger or anticipate chromatin opening^**16-23**^. Thus, GATA factors often occupy a top position within the hierarchies driving differentiation, especially within endoderm lineages. A paradigmatic example is provided by GATA6, which drives the early embryonic segregation of the Primitive Endoderm (PrE) that will lead to extra-embryonic tissues such as the yolk sac, from the pluripotent epiblast – the source of somatic tissues^**24**^.

Rapidly recognized as an essential regulator of the PrE and its first derivatives^**25**^, GATA6 was shown to drive the differentiation of mouse Embryonic Stem (ES) cells, derived from the undifferentiated epiblast, into PrE-like cells^**14**^. *In vivo*, GATA6 was shown to first mark, and then trigger PrE differentiation, via a temporally-ordered circuitry whereby it activates SOX17, a High Mobility Group pioneer TF^**26**^, GATA4 and SOX7, all contributing to efficient PrE differentiation^**13,27-32**^. However, although acting as the upstream regulator in this circuit, GATA6 is unlikely to act alone. Indeed, in line with work suggesting that pluripotency TFs are generally involved in differentiation^**33**^, numerous studies indicate that they contribute to PrE differentiation, suggesting a functional interaction with GATA6 the nature of which and the potential mutual dependencies are not understood. These include SOX2 or NANOG, shown to bind with GATA6 to preserve plasticity and prime PrE differentiation in early differentiating ES cells^**22,34**^; OCT4, a recognized pioneer TF^**35**^ that is required for PrE differentiation *in vivo*^**36-38**^ and promotes PrE differentiation upon its overexpression in ES cells^**39**^ and its interaction with SOX17^**40**^; ESRRB, capable of binding nucleosomal DNA *in vitro*^**41**^, this nuclear receptor first primes^**42,43**^ and then promotes PrE differentiation^**44,45**^ upon the loss of NANOG^**43**^.

Overall, several lines of evidence indicate that the differentiation into PrE requires both the activity of GATA6, potentially as a pioneer TF, and that of several other PrE and pluripotency regulators, which also display pioneering properties. Given the cooperation existing between the pioneer TFs TFAP2A and OCT4^**46**^ to confer developmental competence to neural crest cells^**47,48**^, and the cooption of somatic pioneer TFs by OCT4/SOX2 during cellular reprogramming^**49**^, we aimed at longitudinally exploring how GATA6 induction in ES cells induces transcriptional changes in relation to its pioneering role and the activity of SOX17, OCT4, SOX2 and ESRRB. We found that they engage in a complex network of molecular and epistatic interactions involving additional regulators such as ESRRA, to execute the multitude of gene regulatory tasks required for PrE differentiation, highlighting the large cooperativity that exists between available pioneer TFs during cell fate changes.

## Results

### Distinct dynamics of GATA6 binding are associated with distinct gene regulatory outcomes

To study how GATA6 triggers PrE differentiation we generated ES cells expressing GATA6 under the control of a doxycycline (Dox) inducible promoter^**14,50**^. Upon Dox treatment, ES cells underwent drastic morphological changes over the course of several days **(Fig.S1A)**, accompanied by a progressive modification of marker gene expression from pluripotency (OCT4, SOX2, ESRRB) to PrE-specific proteins (GATA6, SOX17, PDGFRA), characterized by a substantial and transient overlap between the two categories of a priori opposing TFs **(Fig.S1B,C)**. Transcriptomic comparison of undifferentiated ES cells, GATA6-induced cells and XEN cells, a stem cell population derived from early mouse embryos and reminiscent of the PrE^**51**^, further confirmed the progressive nature of the conversion **(Fig.1A)**. We identified 6 clusters of genes up- or downregulated in successive waves **(Fig.S1D)**, initiated either rapidly after Dox induction or at later stages **(Fig.1B and Table S1)**. We then interrogated whether these clusters were enriched in up- or downregulated genes in the PrE of the blastocyst^**52**^. We found that only genes deregulated at early stages displayed a global concordant regulation in the PrE **(Fig.1C)**. Notable examples of early down- and upregulated genes are pluripotency TFs such as *Nanog, Esrrb*, or *Tfcp2l1*, and known drivers of PrE differentiation such as *Sox17, Gata4*, or *Sox7*, respectively **(Table S1)**. We conclude that GATA6 elicits a fast and PrE-specific gene response that rapidly down-regulates pluripotency genes and simultaneously up-regulates PrE genes. To test whether the observed gene responses are linked to GATA6, we performed genome-wide localization studies. We observed global changes in GATA6 binding from 8h of induction to 4 days, with a progressive acquisition of a binding profile similar to that observed in XEN cells **(Fig.1D)**. We identified 3 global behaviors of GATA6 **(Table S2)**, easily observed in single loci **(Fig.S1E,F and Fig.1E)**: regions rapidly targeted by GATA6, which can either maintain (thereafter Early sites) or instead subsequently lose binding as PrE differentiation advances (thereafter Transient sites), or sites that are bound by GATA6 only once differentiation has substantially progressed (thereafter Late sites). To provide support to these dynamics, we compared the binding sites reported here with a fully independent dataset of GATA6 binding in a related population of naïve PrE cells (nEnd^**34**^). As expected, we found a large overlap with Early and Late clusters only **(Fig.1F)**. Upon statistical confrontation^**53**^ of the 3 GATA6 clusters to the 6 groups of differentially expressed genes, we revealed a very clear relationship, with Early sites being strongly enriched near genes that are rapidly activated **(Fig.1G, left panel)** and Late sites being associated with genes that are activated later **(Fig.1G, middle panel)**. In contrast, Transient sites were enriched around genes that are downregulated early or at mid-stages of the differentiation induced by GATA6 **(Fig.1G, right panel)**. Thus, the temporal and dynamic aspects of GATA6 binding are associated with the kinetics (early/late) and modalities (up/down) of gene expression changes driving PrE differentiation.

**Fig. 1.**
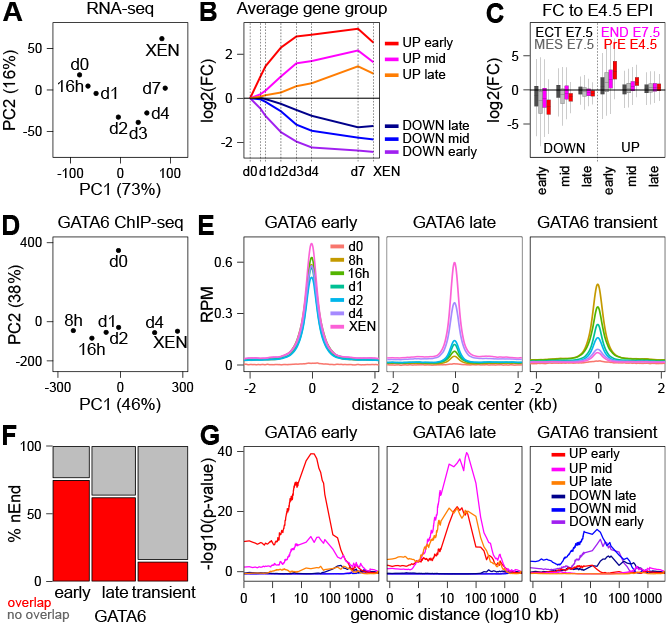
Different dynamics of GATA6 binding induce successive waves of gene expression changes. **(A)** Principal component analysis (PCA) of differentially expressed genes across all analyzed time-points and in XEN cells. **(B)** Average expression profile of 6 distinct clusters of early, mid or late up- or downregulated genes. **(C)** Average log2 fold-change of the 6 gene clusters shown in (B) between the E7.5 ectoderm, endoderm and mesoderm (ECT, END and MES, respectively), or the E4.5 PrE, compared to the E.4.5 epiblast. **(D)** PCA of GATA6 binding regions identified across time-points and in XEN cells. **(E)** Average GATA6 enrichment profiles expressed in reads per millions (RPM) across time-points and in XEN cells for 3 clusters displaying different temporal binding dynamics, centered on the GATA6 summit. **(F)** Proportion of GATA6 binding regions in each category shown in (E) that overlap with binding regions identified in naïve Endoderm cells (nEnd). The statistical significance of the enrichment observed at Early and Late sites was tested with a Chi-square test followed by a Fisher Exact test (p < 2.2e-16). **(G)** Statistical association between each GATA6 binding cluster and the 6 groups of differentially expressed genes shown in (B), measured with Fisher Exact tests at increasing distances.

### Chromatin accessibility and DNA binding motifs underlie GATA6 binding dynamics

We then aimed at identifying the molecular properties that could underlie differential recruitment kinetics and stability of GATA6. We reasoned that the first binding events could be associated with pre-existing chromatin accessibility. To test this, we ranked all GATA6 binding sites by decreasing levels of accessibility in undifferentiated ES cells **(Table S2)** and found a strong correlation with the binding levels observed at early time-points, especially after 8h of Dox induction, for each group of GATA6 binding regions **(Fig.2A)**. Moreover, Late regions were those displaying the lowest proportion of already accessible sites and, conversely, Early and more especially Transient sites displayed a higher proportion **(Fig.2A)**. Even though the enrichment of accessible sites over Transient regions is statistically significant, as is their depletion over Late sites **(Fig.S2A,B,C top panels)**, which are also enriched for heterochromatin marks **(Fig.S2A,B,C bottom panels and Table S2)**, these relationships are not absolute. Thus, additional parameters may contribute to GATA6 dynamics. Accordingly, we found that Early and Late regions displayed the highest and lowest density of high quality GATA6 DNA motifs, respectively **(Fig.2B, Fig.S2A,B,C and Table S2)**. At Transient sites, which display an intermediary average occurrence of GATA6 motifs, as well as at a small proportion of Late sites, we observed a negatively correlated presence of good GATA6 motifs with chromatin accessibility: the less the regions are accessible, the highest is the density and the quality of GATA6 motifs **(Fig.2B)**. This suggests that a combination of pre-existing accessibility and the occurrence of good GATA6 motifs drive, at least partially, GATA6 binding dynamics, in such a way that at highly accessible regions, motifs of less quality are sufficient to trigger fast GATA6 binding and, in reverse, at regions with closed chromatin very good motifs are needed. Inspection of other motifs showed that Transient sites were characterized by a low presence of the PrE TF SOX17 motif and an enrichment of good motifs for pluripotency TFs OCT4/SOX2 and ESRRB **(Table S2)**, especially over accessible regions **(Fig.2B)** bound by pluripotency TF before differentiation **(Fig.2C)**. Conversely, Late sites displayed the opposite pattern **(Fig.2B and Fig.S2A,B,C)** with high quality SOX17 motifs **(Fig.2B)** and an almost complete lack of pluripotency TF motifs **(Fig.2B)** and binding **(Fig.2C)**. Therefore, different combinations of TF motifs likely contribute to differential GATA6 binding dynamics: Early regions are strongly enriched for good GATA6 and SOX17 motifs; Late regions are more strongly associated with SOX17 motifs only; Transient GATA6 binding regions are characterized by the presence of pluripotency TF motifs, coinciding with extensive pluripotency TF binding **(Fig.2C)** and chromatin accessibility **(Fig.2B)**. In support, both DNA motifs and chromatin states, particularly GATA6 motifs and chromatin accessibility, were found as good predictors of the classification of GATA6 binding regions using machine learning **(Fig.S2D)^54^**.

**Fig. 2.**
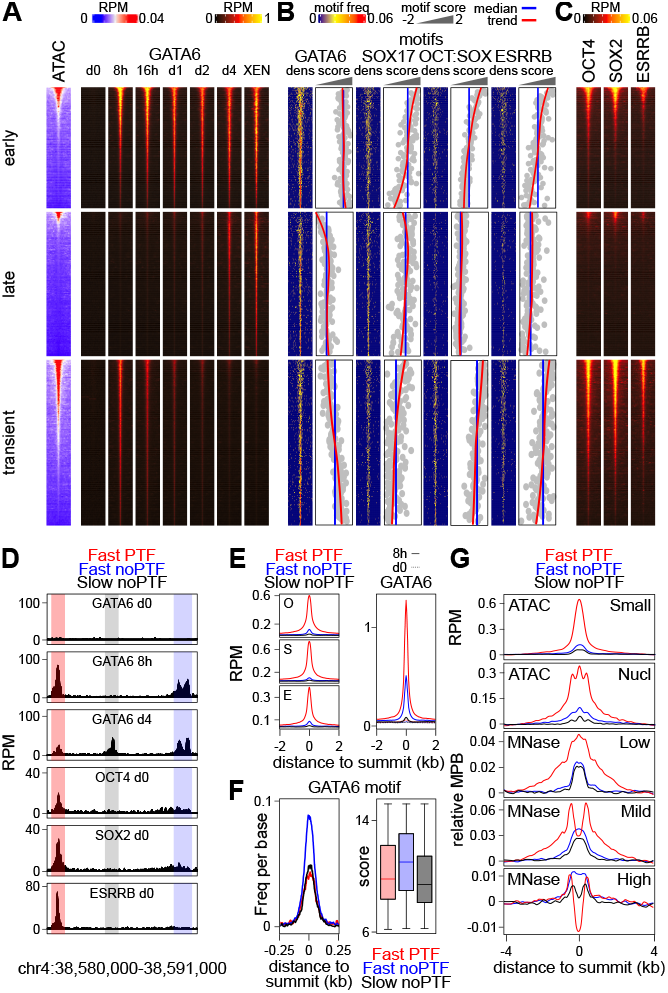
Pre-existing chromatin accessibility, DNA binding motifs and pluripotency TF binding in undifferentiated ES cells, underlie GATA6 binding dynamics. **(A)** Heatmaps depicting chromatin accessibility (ATAC-seq) and GATA6 enrichments, over 4kb-long regions centered on GATA6 summits, ranked by decreasing accessibility in ES cells at each GATA6 cluster. **(B)** Occurrence and quality (identity score to the consensus of the best motif of each region, z-scored to scale between different motifs) of DNA binding motifs across the regions shown in (A), in the same order as in (A). Within each score panel is shown the global median of all regions per cluster (blue) and a trend line (red) of the data (gray). **(C)** Analysis of pluripotency TF binding (reads per million; RPM) across the 3 GATA6 clusters, presented and ordered as in (A). **(D)** Example of different behaviors of GATA6 and pluripotency TFs (PTF). **(E)** Left; average profile of OCT4 (O), SOX2 (S) and ESRRB (E) at the 3 groups of regions illustrated in (D). Right; binding of GATA6 across the same clusters. All 4 plots show the average profile (RPM) over 4kb centered on the GATA6 summit. **(F)** Distribution of GATA6 motifs around the GATA6 summit (left panel), together with the motif score of the best motif present in each region (right panel). Average profiles of chromatin accessibility (small ATAC-seq fragments) and nucleosomes (measured by long fragments detected by ATAC-seq or by MNase-seq using different concentrations of MNase – Low, Mild, High). ATAC-seq is expressed in RPM and MNase-seq in normalized midpoints per billion reads (MPB; see Methods for details). All plots show the average profile over 8kb centered on the GATA6 summit.

### Pluripotency TF binding sites harboring fragile nucleosomes are robustly targeted by GATA6

To more directly explore the relationships between pluripotency TFs, chromatin accessibility and the dynamics of GATA6 binding, we combined Early and Transient regions, which display fast GATA6 recruitment, and divided them as associated with pluripotency TF binding or not **(Fig.2D)**. For comparison purposes we used Late GATA6 regions, which recruit GATA6 very slowly and generally lack pluripotency TF binding **(Fig.2A,D)**. We found that regions of fast GATA6 recruitment displaying pluripotency TF binding in ES cells **(Fig.2E, left panels)** showed the highest levels of GATA6 binding at 8h **(Fig.2E right panel)**, even though these are not the regions with the highest density nor the best quality of GATA6 motifs **(Fig.2F)**. Indeed, the regions displaying the best motif composition are those recruiting GATA6 rapidly, albeit at more modest levels, in the absence of pluripotency TF binding **(Fig.2E,F)**. As expected, regions bound by pluripotency TFs were highly accessible in ES cells compared to those lacking their binding **(Fig.2G, top panel)**, suggesting that their binding might be associated with reduced nucleosomal occupancy. However, when we profiled nucleosomes in the same dataset **(Fig.2G second top panel)**, the only observed difference between the 3 groups was the amplitude of the signal, which represents differential accessibility rather than specific positioning of the nucleosomes. In fact, all 3 regions show the presence of 3 nucleosomes, including a central prominent one located right on the GATA6 binding summit **(Fig.2G second top panel)**. While this is in accord with the pioneering nature of GATA factors^**15,16,23,55**^, the simultaneous detection of accessibility and nucleosomes suggest that they are fragile^**56**^. To test this, we explored previous data generated using increasing concentrations of Microccocal Nuclease (MNase) to map nucleosomes with different stability in ES cells^**57**^. We observed that regions binding pluripotency TFs display a strong nucleosomal signal at the GATA6 summit exclusively when low MNase conditions are used **(Fig.2G; Low)**, with nucleosome signal progressively decaying with increased MNase concentrations **(Fig.2G; Mild)** until it completely disappears at the highest MNase digestion conditions, leading to a profound and apparent nucleosome depleted region **(Fig.2G; High)**. In contrast, at regions lacking pluripotency TF binding the observed nucleosome profiles were less sensitive to MNase digestion conditions **(Fig.2G)**. Since these regions are not accessible, we conclude that they are characterized by more stable nucleosomes. Thus, pluripotency TFs fragilize nucleosomes, leading to a net advantage for fast and robust GATA6 binding even at regions harboring GATA6 motifs of relatively low quality.

### SOX17 recruitment supports long-term GATA6 binding and extensive nucleosome remodeling

Next, we sought to investigate the evolution of chromatin accessibility at Early, Late and Transient GATA6 binding regions **(Table S2)**. We found that at already open Early and Late sites chromatin accessibility was maintained throughout differentiation, whereas at Transient sites it was eventually lost by day 4 of GATA6 induction **(Fig.3A)**. At the remaining and initially closed Early and Late sites, but not at Transient sites, a global strong increase was instated **(Fig.3A)**, rapidly at Early sites (by days 1/2) and more slowly at Late sites (by days 2/4). However, these dynamics were relatively delayed compared to the timing of GATA6 binding over these groups, especially at Early sites that bind GATA6 as early as 8h following its induction **(Figs.1,2)**. Given the prevalence of SOX17 DNA motifs over Early and Late regions compared to Transient sites **(Fig.2B)**, and the fact that GATA6 binds rapidly at the *Sox17* locus to activate its expression **(Fig.S3A,B and Fig.S1C)**, we reasoned that it could contribute to chromatin opening of these sites. Hence, we profiled SOX17 binding and found that it is efficiently recruited to a large fraction of GATA6 bound regions **(Fig.3B, Fig.S3B,C and Table S2)**, predominantly at Early and Late sites and only anecdotally at Transient sites **(Fig.3B, Fig.S3C)**, matching the distribution of its DNA motif **(Fig.2B, Fig.S2)**. Moreover, comparing the average trends of chromatin accessibility, GATA6 and SOX17 recruitment, we observed that SOX17 is recruited concomitantly to, or slightly before, any measurable increase in chromatin accessibility, whereas GATA6 binding occurs earlier **(Fig.3C)**. Accordingly, when we separately analyzed the regions as a function of SOX17 binding, we confirmed that those recruiting SOX17 tend to recruit higher levels of GATA6 and instate more efficiently chromatin accessibility **(Fig.S3D,E)**. Therefore, SOX17, an early target of GATA6, reinforces GATA6 binding and strongly contributes to chromatin opening. Motivated by these observations, we aimed at analyzing nucleosome profiles. For this, we first used our accessibility datasets, which revealed the presence of 3 well positioned nucleosomes over the analyzed regions, without making any distinction between GATA6 binding groups beyond the reported changes in global accessibility **(Fig.S3F)**. Thus, we performed MNase digestions using sufficiently drastic conditions to observe the disappearance of fragile nucleosomes, as established above **(Fig.2G)**. For these analyses, and with the aim of avoiding confounding variables due to the presence of distinct initial states driven by pluripotency TFs **(Fig.2)**, we separated GATA6 binding regions by their dynamics (Early, Late, Transient), by the presence of pluripotency TFs before differentiation and by the recruitment of SOX17 during differentiation, excluding combinations that were poorly represented **(Fig.S3C, Table S2)**. As expected, only sites with pluripotency TF binding displayed an apparent central nucleosome depleted region before GATA6 induction **(Fig.3D)**, testifying of the presence of fragile nucleosomes. Upon GATA6 induction, the profiles changed drastically, especially over regions recruiting SOX17, more rapidly at Early than at Late sites and only transiently at Transient sites **(Fig.3D)**. By day 4 of differentiation, the initial differences mediated by pluripotency TFs had been fully erased, Transient sites showed no signs of fragile nucleosomes (in accord with reduced levels of accessibility), and Early and Late sites showed precisely ordered nucleosomal arrays flanking an apparent nucleosome depleted region at the GATA6 summit, a profile that was more pronounced over regions also bound by SOX17 **(Fig.3D)**. Assessment of nucleosome order over time and across GATA6 binding groups **(Fig.3E)** provided quantitative support to the conclusion that GATA6 initiates, and SOX17 drastically enforces, an extensive nucleosome remodeling that leads to properly ordered nucleosomes around a central fragile nucleosome sitting at the GATA6 binding summit.

**Fig. 3.**
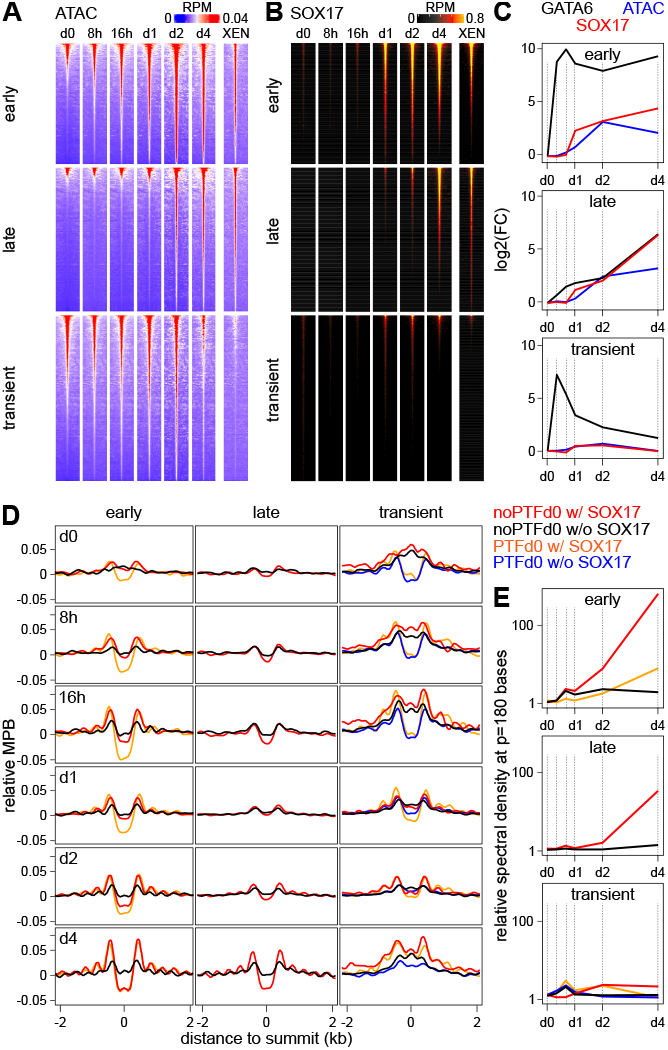
SOX17 recruitment promotes and stabilizes GATA6 binding leading to increased chromatin remodeling. **(A)** Evolution of chromatin accessibility measured by ATAC-seq over GATA6 binding groups. Presented and ordered as in Fig.2A. **(B)** Evolution of SOX17 binding across the same regions, ordered as in A and Fig.2. **(C)** Average log2 Fold-Change of chromatin accessibility, GATA6 and SOX17 binding at every time-point versus undifferentiated cells (d0). **(D)** Evolution of nucleosome positioning at the most prominent GATA6 binding categories shown in Fig.S3C, presented as in Fig.2G. **(E)** Quantification of nucleosome order using the spectral density calculated from the profiles shown in (D) at a period of 180 bases – nucleosome (150 bases) plus linker DNA (30 bases) – throughout GATA6 induction time-points.

### OCT4/SOX2 promote nucleosome remodeling at GATA6/SOX17 targets and PrE gene activation

We next aimed at monitoring pluripotency TF binding along GATA6 induction, with the initial idea of following how the pluripotency network is dismantled. Upon OCT4, SOX2 and ESRRB profiling **(Fig.4A, Fig.S4A and Table S2)**, we observed that GATA6 binds consistently to around a fifth of all sites bound by pluripotency TFs in undifferentiated ES cells **(Fig.4A)**. Moreover, GATA6 preferentially targets regions characterized by high levels of OCT4 and SOX2 binding, especially when ESRRB is also bound **(Fig.4A and Fig.S4B)**. In contrast to our expectations, we found that at regions targeted by GATA6, binding of pluripotency TFs was detected during a longer period after GATA6 induction than at regions not targeted by GATA6, which lose almost all signs of pluripotency TF binding during the very first days of GATA6 induction **(Fig.4A and Fig.S4B)**. While a global reduction of pluripotency TF binding is expected given the downregulation of their expression **(Fig.S1B,C and Table S1)**, their transient retention at GATA6 bound regions indicates that the declining pool of remaining molecules is specifically redirected at these sites. In addition, our analyses identified a cluster of GATA6 regions experiencing ectopic binding of pluripotency TFs during differentiation, especially of OCT4/SOX2 **(Fig.4A and Fig.S4A,B)**. These regions were found enriched for Early GATA6 binding regions and for SOX17 recruitment **(Fig.4B,C)**, displayed high levels of GATA6 and SOX17 binding and rapidly gained accessibility **(Fig.4D top)** to levels even higher than those observed at the best pluripotency TF sites in ES cells **(Fig.S4C, top row)**. In contrast, pluripotency TF binding sites not targeted by GATA6 progressively lost chromatin accessibility **(Fig.S4C, top row)**. Analysis of nucleosome profiles from accessibility datasets confirmed previous observations, with the detection of nucleosomes at the GATA6 summit **(Fig.S4C middle row)**. Using MNase, we could however unmask the appearance of an apparent nucleosome depleted region concomitantly with ordered flanking nucleosomes **(Fig.4D bottom, Fig.S4C bottom row)**, which was much more pronounced at regions of ectopic pluripotency TF binding than at those not associated with their recruitment. Similarly, focusing on regions bound by pluripotency TFs in undifferentiated ES cells and comparing those that are either targeted or not by GATA6, we could confirm that only the later exhibit a progressive invasion of the region by stable nucleosomes, whereas those associated with GATA6 preserve an apparent nucleosome depleted region that represent the presence of fragile nucleosomes, particularly at early time-points after Dox induction **(Fig.S4C bottom row)**. Furthermore, we found that GATA6 sites with ectopic OCT4/SOX2 binding were strongly and very specifically associated with gene activation events occurring early following GATA6 induction, whereas regions not associated with pluripotent TFs were equally associated with gene activation at any time **(Fig. 4E)**. In contrast, the regions displaying pluripotency TFs before differentiation tend to be associated with early repressive gene changes **(Fig.4E)**. We conclude that GATA6 and SOX17 maintain and repurpose pluripotency TFs binding, leading to a prominent nucleosome remodeling and the rapid activation of PrE genes.

**Fig. 4.**
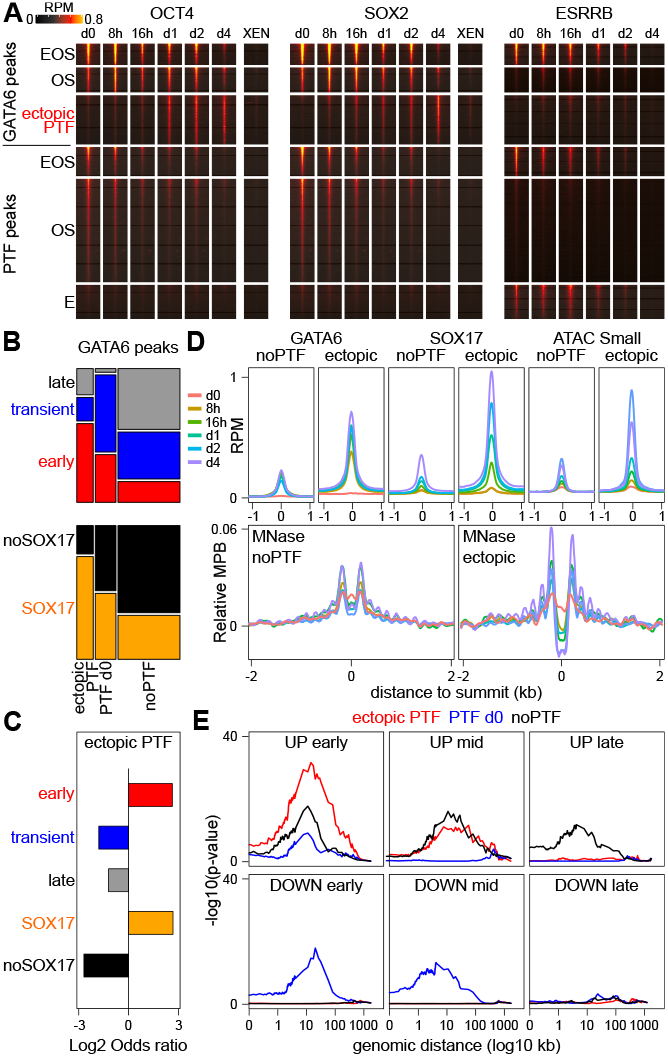
GATA6 maintains and promotes ectopic binding of pluripotency TFs during PrE differentiation. **(A)** Binding dynamics of OCT4, SOX2 and ESRRB upon GATA6 induction, both at GATA6 peaks showing pluripotency TF binding before or during differentiation (i.e. GATA6 peaks lacking pluripotency TF binding are omitted) and at other pluripotency TF binding sites in undifferentiated ES cells. **(B)** Correlations between categories of GATA6 binding segregated by pluripotency TF binding (X-axis) and by GATA6 dynamics (top; Y-axis) or by SOX17 binding (bottom; Y-axis). The dependency between variables was assessed with Chi-square tests (p < 2.2e-16). **(C)** Log2 odds ratio of the proportion of regions displaying ectopic pluripotency TF binding across different GATA6 binding groups shown on the left. The statistical significance of the enrichments/depletions were assessed with Fisher Exact tests (p < 1.3e-155). **(D)** Average profiles of GATA6 binding, SOX17 binding, chromatin accessibility (small ATAC-seq fragments) and nucleosome positioning (MNase-seq) at regions displaying no pluripotency TF binding at all (noPTF) or an ectopic recruitment (ectopic), throughout GATA6 induction (colored lines). All plots are centered on the GATA6 summit. **(E)** Statistical association of GATA6 binding regions divided by pluripotency TF binding groups, with the 6 groups of differentially expressed genes shown in Fig.1B, measured with Fisher Exact tests at increasing distances.

### OCT4 is required for *Gata6* upregulation and PrE differentiation

Given the strong link existing between ectopic OCT4/SOX2 binding and the activation of early GATA6-responsive genes, we decided to test whether OCT4 is strictly required for PrE differentiation. To address this, we resorted to an established chemical induction of PrE differentiation^**58,59**^ and used pluripotent cells that express OCT4 fused to an Auxin inducible degradation domain enabling fast and acute depletion upon IAA treatment^**60,61**^. While IAA-untreated cells efficiently differentiated into PrE **(Fig. 5A,B,C, Fig.S5A,B)**, depleting OCT4 at different time-points had strong and variable consequences. When OCT4 depletion was triggered concomitantly with the onset or at day 1 of differentiation, we observed a drastic mortality of the cells, uncapable of surviving the chemical induction **(Fig.5A)**. In contrast, when OCT4 depletion was performed at day 2 or later during the chemical induction, the differentiation into PrE was as efficient as in IAA-untreated cells **(Fig.5A, Fig.S5A,B)**. Hence, OCT4 is essential to allow early differentiating cells to survive, supporting the notion that the molecular effects of OCT4 at GATA6 targets are necessary for PrE differentiation. Furthermore, the quantification of PrE markers during the chemical induction of differentiation revealed low GATA6 **(Fig.5B,C)** and PDGFRA expression **(Fig.S5A,B)** in the rare cells that succeeded to differentiate upon IAA treatment from day 1, but also in cells experiencing OCT4 depletion from day 2 and even day 3. Exploration of our TF binding data around the *Gata6* locus **(Fig.5D)** identified a region of fast GATA6 and SOX17 binding, where OCT4/SOX2 are ectopically recruited upon GATA6 induction and, notably, which displays the chromatin signature of an active enhancer, as observed using published datasets^**34**^. This suggests that the early effects of OCT4 depletion observed upon chemical induction may also involve the incapacity of the cells to properly induce GATA6 and initiate the large changes required for PrE differentiation. In contrast, once GATA6 has already irreversibly initiated differentiation, the depletion of OCT4 at later stages and the ensuing downregulation of *Gata6* is largely inconsequential.

**Fig. 5.**
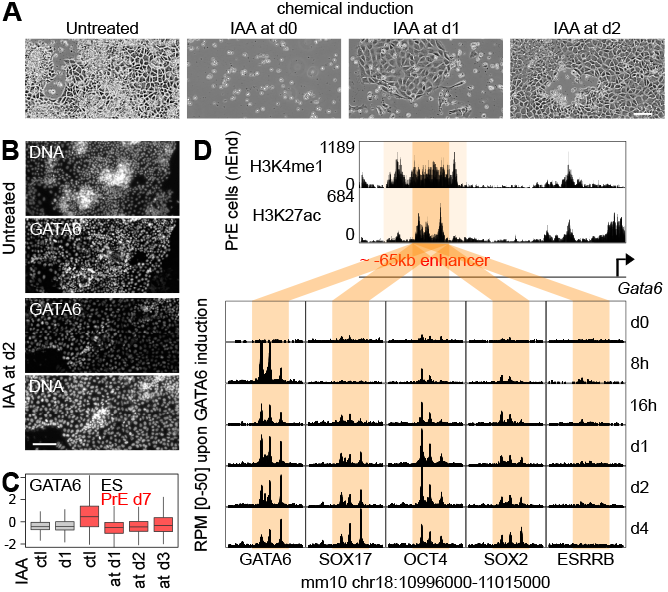
OCT4 is essential during early stages of PrE differentiation. **(A)** Photomicrographs of cells subject to chemical PrE induction for 7 days, either untreated or IAA-treated cells to deplete OCT4 from the onset of the chemical PrE induction (d0), from day 1 (d1) or day 2 (d2). The PrE-like cells shown for d1-treated cells represent one of the rare colonies that did not die. The scale bar represents 100 µm. **(B)**.Immuno-staining of GATA6 after 7 days of chemical PrE induction, for untreated (top) and IAA-treated cells from day 2 onwards (bottom). The scale bar represents 150 µm. **(C)** Quantification (Z-score) of GATA6 immuno-staining in control ES cells (ctl in gray), after 1 day of IAA treatment (d1 in gray), and in cells differentiated for 7 days (in red) in the absence of IAA (ctl) or having experienced OCT4 depletion from day 1, day 2 or day 3 onwards. **(D)** Identification of an active enhancer based on histone marks (top) *∼*65kb upstream of *Gata6* showing no TF binding before differentiation but rapidly recruiting GATA6 and SOX17 and displaying ectopic OCT4/SOX2 binding during GATA6 induction.

### Distinct nucleosome remodeling regimes at ESRRB binding sites predict a role for ESRRA

We have shown that OCT4/SOX2 binding regions that are not targeted by GATA6 display a loss of accessibility and an invasion of nucleosomes during differentiation regardless of ESRRB binding **(Fig.S4C)**, as clearly illustrated comparing uninduced and Dox-induced cells for 4 days **(Fig.6A)**. In contrast, regions bound by ESRRB alone did not display such extensive remodeling **(Fig.6A and Fig.S4C)**, despite the very efficiently downregulation of this TF upon GATA6 induction **(Fig.S1C, Fig.6B)**. This indicates that the regions bound by ESRRB in ES cells might be targeted by additional TFs, responsible for preserving nucleosome order once ESRRB is fully lost. Given the importance of ESRRB during the early stages of PrE differentiation^**42-45**^, we sought to identify the factors that may replace it. ESRRB belongs to the large family of nuclear receptors^**62**^, and thus shares an evolutionary origin and structural properties with related TFs, with which it can exert redundant functions in ES cells as shown for NR5A2 and ESRRA^**63,64**^. Thus, we hypothesized that other nuclear receptors could take over ESRRB function during the induction of GATA6. We probed nuclear receptor expression during GATA6 induction and observed that only 3 showed stable or increased expression, among which only the closely related TF ESRRA is upregulated **(Fig.6B)**. Moreover, ESRRA displays an almost identical binding motif to ESRRB **(inset in Fig.6B)** and, in the blastocyst, shows maximal expression in the PrE compared to other early lineages^**62**^. Thus, ESRRA may functionally replace ESRRB at a subset of its targets, maintaining their activity. In agreement, we found that regions targeted by ESRRB (but not by OCT4/SOX2) in ES cells, are also bound by ESRRA both before and after 4 days of GATA6 induction **(Fig.6C)**. This suggests that ESRRA replaces ESRRB as a nucleosome organizer at a subset of regulatory elements that remain active throughout the whole conversion of ES into PrE cells driven by GATA6. We also found that a reduced proportion of GATA6-bound regions displayed binding of ESRRA before GATA6 induction, largely coinciding with ESRRB-bound regions **(Fig.S6A)**; in contrast, 4 days after inducing GATA6 we observed a larger and more intense ESRRA binding activity at GATA6 targets, despite the loss of ESRRB **(Fig.S6A)**. Regions displaying high levels of ESRRA binding at d4 are enriched in Early GATA6 sites and are associated with SOX17 and ectopic OCT4/SOX2 recruitment **(Fig.S6B**). In line with this, when we separated all GATA6 binding regions in all combinations made by GATA6 dynamics, SOX17 and OCT4/SOX2 binding **(Fig.S6C)**, and focused on the most prominent ones **(Fig.S6C)**, we observed that ESRRA was preferentially recruited at GATA6 binding regions recruiting SOX17, irrespective of ESRRA binding status before differentiation **(Fig.6D, Fig.S6D)**. These regions recruiting SOX17 and ESRRA during differentiation are, especially in combination with ectopic OCT4/SOX2 binding, those most strongly associated with early PrE gene activation **(Fig.6E)**. Thus, not only ESRRA replaces ESRRB during PrE differentiation at a subset of the regions it targets in ES cells, ESRRA also engages at sites targeted by GATA6, in cooperation with SOX17, OCT4 and SOX2, where ESRRB is barely involved.

**Fig. 6.**
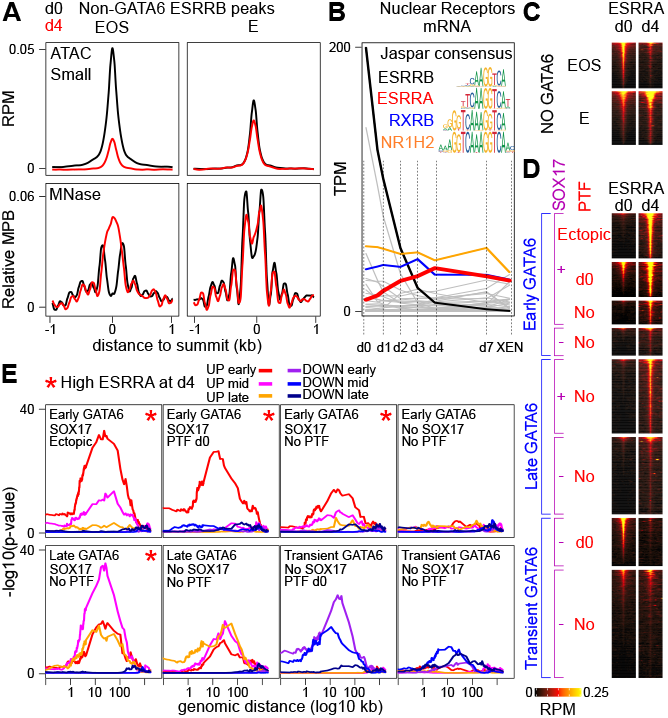
ESRRA replaces and expands ESRRB function during PrE differentiation. **(A)** Average profiles of chromatin accessibility measured by small ATAC-seq fragments (top; Reads Per Million – RPM), together with the average nucleosome positioning profiles measured by MNase-seq (bottom; normalized Midpoints Per Million of nucleosome-sized fragments) at pluripotency TF binding regions that do not recruit GATA6 and that are enriched for either ESRRB, OCT4 and SOX2 (EOS) or exclusively ESRRB (E), before or after 4 days of GATA6 induction. All the plots, extracted from Fig.S4C, are centered on the accessibility summit. **(B)** Expression of all nuclear receptors mRNAs detected by RNA-seq during GATA6 induction (gray light lines), with 4 highlighted in color: *Esrrb, Esrra, Rxrb* and *Nr1h2*. The inset shows the consensus motif of the 4 nuclear receptors from the Jaspar database.**(C)** Enrichment of ESRRA across ESRRB binding sites identified in ES cells and not associated with GATA6 binding, separated depending on whether they recruit other pluripotency TFs (EOS) or exclusively ESRRB (E) after 4 days of GATA6 induction. **(D)** Enrichment of ESRRA across the most prominent GATA6 clusters, as established in Fig.S6C, before and after 4 days of GATA6 induction. **(E)** Statistical association of the groups of regions shown in (D) with the 6 clusters of differentially expressed genes shown in Fig.1B, measured with Fisher Exact tests at increasing distances. The red asterisk denotes clusters with prominent ESRRA enrichment at d4 of differentiation, as shown in (D) and in Fig.S6C,D.

### ESRRA sustains *Oct4* expression and is essential for PrE differentiation

To test whether ESRRA would be, like ESRRB^**42,43**^, important for PrE differentiation, we generated *Esrra* knock-out cells ΔEa cells, **Fig.S7A**) and chemically induced their differentiation into PrE. While wild-type cells efficiently differentiated, ΔEa cells displayed high mortality and almost completely failed in generating PrE cells **(Fig.7A,B,C)**, in a manner similar to that observed during the premature depletion of OCT4 **(Fig.5)**. Surviving ΔEa cells did however exist, and preserved expression of some pluripotency markers, such as NANOG, albeit at low levels **(Fig.7C and Fig.S7B)**.

**Fig. 7.**
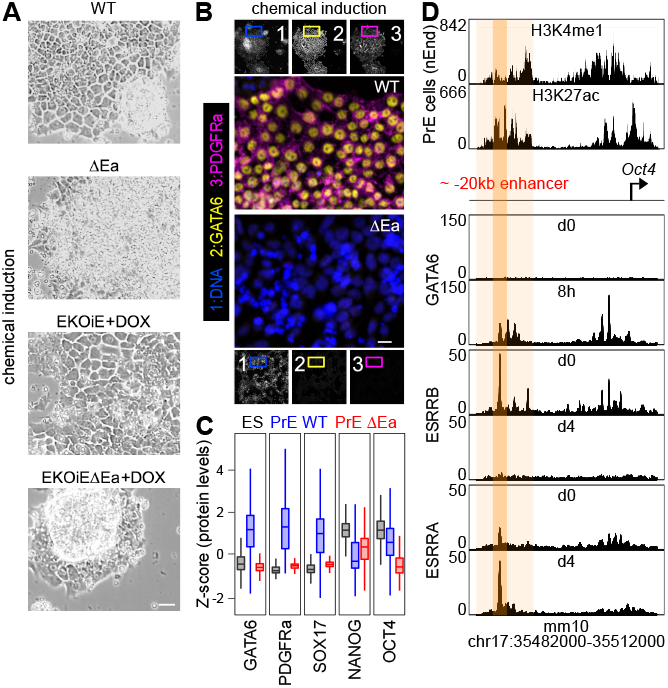
*Esrra*-/- ES cells cannot differentiate into PrE. **(A)** Representative photomicrograph of wild-type (WT) and *Esrra*-/- cells (ΔEa), as well as of cells with forced ESRRB expression (EKOiE and EKOiEΔEa), after 7d of chemical PrE induction. Note the extensive mortality in ΔEa cells. The scale bar represents 50 µm. **(B)** Representative immuno-fluorescence of GATA6 and PDGFRA after 7d of PrE chemical induction. The scale bar represents 20 µm. **(C)** Marker quantification in undifferentiated WT ES cells (black) and in WT and ΔEa cells after 7d of chemical PrE induction (blue and red, respectively). **(D)** Identification of a GATA6-bound active enhancer based on histone marks (top) located 20kb upstream of *Oct4* and showing the replacement of ESRRB by ESRRA upon GATA6 induction.

In contrast, ΔEa cells displayed lower levels of OCT4 than WT cells **(Fig.7C)**, suggesting that ESRRA activates *Oct4* and enables the slow rather than fast decay of OCT4 levels **(Fig.S1B)**, required for its persistent activity during differentiation. Exploration of ESRRA binding in the vicinity of *Oct4* allowed us to identify an active enhancer located *∼* 20kb upstream of its promoter that is targeted by ESRRB in undifferentiated cells and by ESRRA upon GATA6 induction **(Fig.7D)**, providing direct support to the notion that these two nuclear receptors exchange to sustain *Oct4* expression. Next, we asked whether ESRRB could rescue ESRRA deficiency using available ES cells where ESRRB expression is fully controlled by Dox (EKOiE^**65**^). First, we confirmed that ESRRB promotes PrE differentiation^**43**^, since we observed a large and scaled detrimental effect when ESRRB was experimentally silenced during, only before or both before and during the chemical induction of differentiation **(Fig.S7C)**, compared to cells with forced ESRRB expression that differentiated very efficiently **(Fig.S7C and Fig.7A bottom panels)**. Thus, we generated the *Esrra* knock-out in EKOiE cells **(Fig.S7A)** and found that regardless of ESRRB expression, EKOiEΔEa displayed extremely poor PrE differentiation efficiency **(Fig.7A bottom panels and Fig.S7C)**. We conclude that GATA6 promotes ESRRA expression, which replaces ESRRB and extends its function to a point that it cannot be complemented by ESRRB, by targeting regions bound by GATA6, including an *Oct4* enhancer.

## Conclusion

### Molecular roles and epistatic interactions of GATA6 during PrE differentiation

GATA6 leads to PrE differentiation via 3 major waves of gene regulation, with the earliest ones initiating the downregulation of the players of the pluripotency network and the up-regulation of PrE genes, including SOX17. These regulatory waves can be ascribed to specific modalities of GATA6 binding among its 8 main categories **(Fig.8)** and are supported by extensive remodeling of the gene regulatory network (see epistatic diagram in **Fig.8**). Opportunistic GATA6 binding at regions made accessible by pluripotency TFs is fast even if GATA6 motifs are not particularly good, thanks to the fragilization of nucleosomes mediated by OCT4/SOX2/ESRRB. At these sites, good motifs for SOX17 and its subsequent recruitment determine if GATA6 binding will be stable or, on the contrary, transient. While transient GATA6 binding at these regions leads to their decommissioning and the dismantlement of pluripotency, stable GATA6/SOX17 binding leads to early PrE gene activation. GATA6 also binds at closed regions, acting as a pioneer TF with rapid or slow dynamics depending of the quality of GATA6 and SOX17 motifs. In the presence of good GATA6 motifs, its binding is fast and stable but, without the intervention of other TFs, it does not lead neither to strong nucleosome remodeling nor to particular gene responses. In contrast, when GATA6 is accompanied by SOX17, pluripotency TFs transiently repurposed for differentiation (OCT4/SOX2), and the nuclear receptor ESRRA, it leads to drastic nucleosome remodeling and early activation of PrE genes. When GATA6 motifs are of poorer quality, but provided that good SOX17 motifs are present, GATA6, SOX17 and to a lesser extent ESRRA, are recruited later, leading to late gene activation. Thus, while GATA6 initiates a profound nucleosome remodeling characterized by the acquisition of central fragile nucleosomes flanked by ordered nucleosomal arrays and increased chromatin accessibility, this is fully maximized only upon binding of SOX17, OCT4, SOX2 and/or ESRRA. These TFs, which are regulated by GATA6, are essential for PrE differentiation and positively feed-back on *Gata6* via a self-reinforcing *Gata6-Esrra-Oct4-Gata6* loop. Thus, we have identified different modes of direct and indirect action of GATA6 to activate or repress transcription, acting either as an opportunistic or a pioneer TF and unfolding epistatic interactions to rewire the gene regulatory network **(Fig.8)**.

**Fig. 8.**
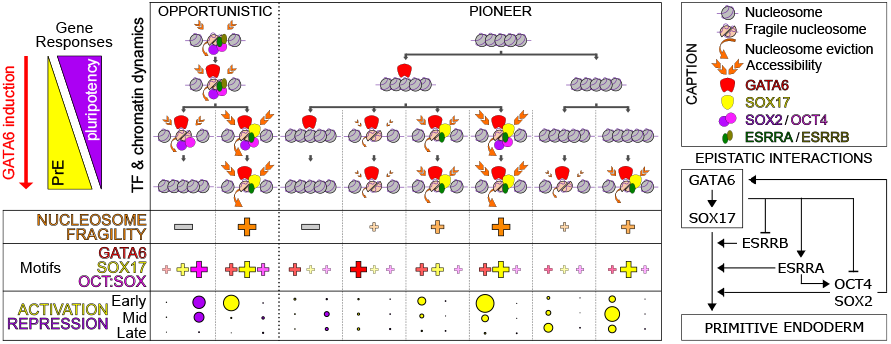
Direct and indirect functional interactions mediated by GATA6. Summary of the molecular observations reported in this study, highlighting different modes of GATA6 binding (opportunistic or pioneer), different dynamics (early versus late and stable versus transient) and different outcomes regarding: ectopic recruitment of SOX17, OCT4/SOX2 and ESRRA; chromatin structure; gene regulatory influences at early, mid or late responsive genes; the occurrence of DNA motifs. A caption and a diagram of the epistatic interactions emerging from kinetic analyses and knock-out cells are shown.

## Discussion

### GATA6, an example of cooperative action within a “pioneerosome”

This study adds to the broad literature supporting a role for GATA factors in pioneering processes^**10,11,15–23**^. However, we show here that GATA6 maximizes its activity in the context of a “pioneerosome”, defined as the cooperative action of several pioneer TFs at defined regulatory elements. This notion might be particularly relevant for GATA factors, for which their pioneering activity is, to some extent, surprising, given that they have two consecutive zinc fingers that bind high affinity motifs in free DNA but pose steric hindrance issues when DNA is bent in a nucleosome. For GATA3, like for other pioneer TFs binding partial motifs embedded in nucleosomes^**26,35,55,66,67**^, this may be alleviated thanks to its nucleosomal binding to specific motif geometries consisting of repeated partial motifs^**16**^. However, we observe a strong dependence of GATA6 to its cognate motif, especially at inaccessible sites. As suggested for SOX2^**68**^, this could depend on the high GATA6 levels attained in our experimental system. However, GATA6 motifs were already found of better quality in nucleosomal than in free DNA in an independent study that did not use overexpression^**55**^. Strikingly, these good GATA6 motifs were present in regions also targeted by HNF3^**55**^, which exhibits greater pioneering capacity than another GATA factor, GATA4^**15**^. This suggests that cooperative nucleosome attack by several pioneer TFs may help recognition and binding to canonical motifs even when embedded in nucleosomes. Accordingly, GATA6 deploys proper pioneering activity in the presence of other pioneer TFs such as OCT4, SOX2 and SOX17. Thus, our data calls for a more nuanced view on the pioneering nature of GATA6, and, by extension, of other GATA factors. As proposed for SOX2^**69**^, GATA6 acts both as a “settler” and opportunistic TF, when it binds at open regions and decommissions pluripotency enhancers, or as a “pilot” TF guiding further pioneer TF recruitment and remodeling activities **(Fig.8)**. Of course, we show that GATA6 binds at nucleosomal DNA, regardless of whether it is already made accessible by OCT4/SOX2 or if it is located in closed chromatin, be it naïve or heterochromatic. This, by itself, qualifies GATA6 as a pioneer TF; it remains true, however, that its binding is faster and more robust, and its effects on nucleosomes and accessibility greater, when occurring in cooperation with other TFs. Even though GATA6 effects on chromatin manifest as soon as it binds, one to two days are needed to fully fragilize the nucleosome it targets, position the flanking ones as an ordered array, and fully open up the chromatin. This latency could be dependent on the need for DNA replication, which disrupts nucleosomes^**70**^ even when targeted by pioneer TFs^**71**^, but its correlation with the recruitment of other pioneer TFs indicates that GATA6 initiates a process that is fully culminated by cooperative activities. Hence, cooperative interactions between pioneer TFs and the formation of a “pioneerosome” appear instrumental in the way GATA6 unfolds the gene expression changes that lead to PrE differentiation.

### GATA6 rewires the gene regulatory network

The role of *Oct4* in the establishment of the PrE was suggested more than 20 years ago^**36,39**^. Even though it has been debated whether its role is cell autonomous^**37**^ or not^**38**^, OCT4 contribution to PrE differentiation, while clear, had remained mechanistically enigmatic until now. With this study, we propose that OCT4 is both an activator of *Gata6* and a mediator of its early effects, since it is redirected by GATA6 to readily enact differentiation. This scenario, strictly reciprocal to the redistribution of somatic TFs upon OCT4/SOX2-induced reprogramming back to pluripotency^**49**^, is different than previous propositions for a role of pluripotency TFs in priming, extending or restoring developmental potential during lineage commitment^**22,34,46–48,72**^. Moreover, our observation that in the early absence of OCT4 during PrE differentiation, cell viability is compromised, opens non-mutually exclusive interpretations of its relevance. On the one hand, it is possible that GATA6-OCT4/SOX2 activate expression of genes directly involved in cell survival – anti-apoptotic genes (*Bcl2l1*/*Bcl-xl, Birc5*/*Survivin* and *Birc6*/*Bruce*) and genes with a role in PrE survival^**73,74**^ (*Pdgfra, Akt1* and *Akt2*) are upregulated upon GATA6 induction **(Table S1)**, with PDGFRA expression displaying an attenuated increase in OCT4-depleted cells. On the other, the early loss of OCT4 may lead to a molecular context of inadaptation between the signals pushing towards the PrE, the propensity of *Oct4*-/- cells to differentiate along the trophectoderm^**36,39**^, and their failure to activate *Gata6*, resulting in cell death. The observation that the lack of ESRRA, which is required for sustained OCT4 expression during early PrE differentiation, also leads to increased cell death lands support to this interpretation. ESRRA is upregulated upon GATA6 induction and replaces ESRRB, a related TF that is rapidly turned off despite its role as a PrE priming TF^**42-45**^. Hence, through the concomitant compensation of ESRRB downregulation by ESRRA up-regulation, GATA6 coordinates a switch between nuclear receptors to sustain *Oct4* expression, maintain accessible those regulatory elements that were specifically controlled by ESRRB in ES cells, and broadly extend their function to regions targeted by GATA6. While the functions of nuclear receptors are redundant in pluripotent cells^**63,64**^, this and other studies indicate that their importance can also be hierarchized as changes in cellular contexts are implemented^**75**^. Whether the essential role of ESRRA is exclusively mediated by its direct enhancement of GATA6 activity or also indirectly via the control of *Oct4*, or whether it is important for other regulations needed for differentiation and survival^**76-78**^, such as mitochondrial activity and lipid catabolism^**79**^, remains unclear. While it is likely that GATA6 activates other TFs that will exert additional functions, these observations indicate that this pioneer TF unfolds a large regulatory network that goes beyond its immediate molecular role.

In summary, we show here that GATA6 takes over a multitude of gene regulatory processes, taking advantage of both the *cis*-landscape of DNA binding motifs and pre-established chromatin states, as well as the *trans*-compendium of available pioneer TFs. This enables GATA6 to multitask and to organize its activity over time, such that it can repress pluripotency, timely initiate pioneering activities at new regulatory elements required for successive waves of PrE gene activation and promote new TFs to higher hierarchies to extend its effects beyond its direct binding sites. Thus, even though pioneer TFs are especially attractive for their unique capacity in licensing close regulatory elements^**80**^, their importance in developmental regulation goes beyond this key structural property. They fully rewire gene regulatory networks to endow the transcriptional changes required to acquire new cell identities.

## Supplementary information

7 supplementary figures can be found at the end of this document and 2 Supplementary Tables and Methods online.

## Acknowledgements

The authors acknowledge Jose Silva and Lawrence Bates for the kind gift of *Oct4*-AID cells, all members of the laboratory for critical discussions and the OMICs facility of Institut Pasteur for access to sequencing platforms. This study was funded by the Labex Revive (ANR-10-LABX-73; R.X.C, M.C.T, P.N.), the CNRS (M.C.T and P.N.), the Institut Pasteur (P.N.), the ANR (ANR-14CE11-0017PrEpiSpe; M.C.T.) and the European Research Council (ERC-CoG-2017 BIND; P.N.)

## Contributions

This study was conceived by P.N. and M.C.T. Experiments were designed and executed by R.X.C. and A.D. (chemical PrE induction and ESRRA ChIP-seq), with help from I.G. and S.V.P. *Esrra* knockout cells were generated by N.F. and characterized by A.D. Gene expression analyses were performed by A.C. and P.N.; chromatin analyses were performed by R.X.C and P.N. The paper was written by R.X.C, M.C.T. and P.N. and approved by all authors.

## Declaration of interests

The authors declare no competing interests.

---

**Supplementary Information, Fig. S 1.**
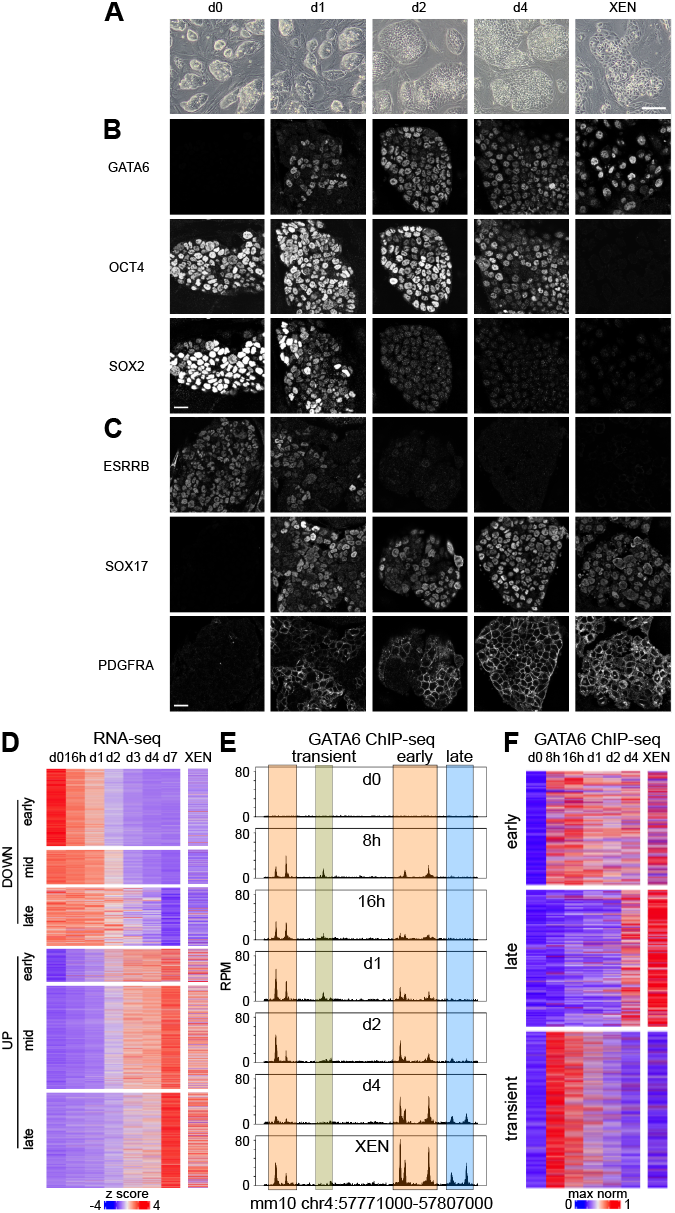
Conversion of ES cells into PrE cells by GATA6 induction. **(A)** Illustrative photomicrograph of ES cells (d0) and as they convert into PrE cells following Dox treatment for 1, 2 and 4 days, as well as of XEN cells. The scale represents 200 µm. **(B)** Representative immunostaining of GATA6, OCT4 and SOX2 in the same samples as in (A). The scale represents 30 µm. **(C)** Representative immunostaining of ESRRB, SOX17 and PDGFRA in the same samples as in (A). The scale represents 30 µm. **(D)** Expression changes of the 6 groups of genes identified as early/mid/late up- or downregulated. **(E)** Illustrative locus showing different behaviors of GATA6 binding dynamics. **(F)** Relative GATA6 binding levels across the 3 clusters of GATA6 binding regions. For each region, the sample with maximal enrichment levels was set to 1.

**Supplementary Information, Fig. S 2.**
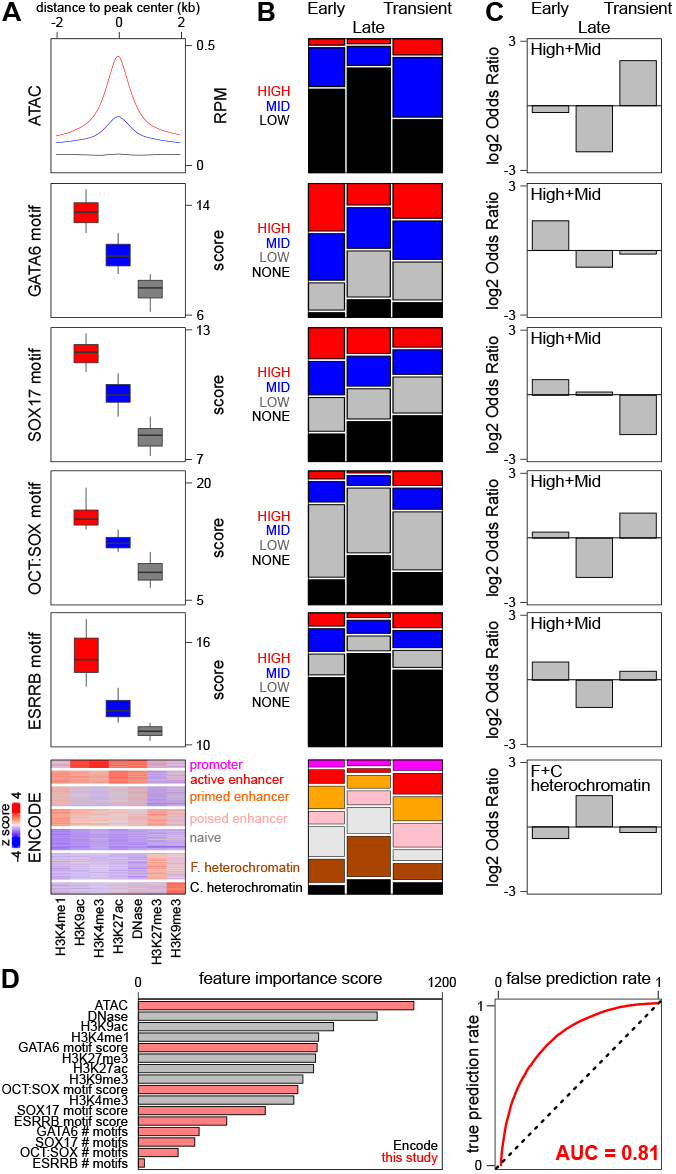
Statistical analyses of GATA6 binding regions. **(A)** Several properties are considered and the regions divided in 3 or more groups, from top to bottom: accessibility measured by small ATAC-seq fragments, with the regions divided as displaying high, mid or low accessibility; the quality of the best motifs of GATA6, SOX17, OCT:SOX or ESRRB present in each region measured by their score, with the regions divided as presenting motifs with high, mid, low scores or none at all; histone modification status as evaluated by quantification and clustering of Encode datasets, grouping the regions according to the presence of marks typical of distinct regulatory states, shown in the right (F. facultative; C. constitutive). **(B)** Proportion of the different groups of variables shown in (A) throughout Early, Late and Transient GATA6 binding regions. The statistical dependence between each group of variables and the 3 groups of GATA6 binding regions was shown to be significant using Chi-square tests for all combinations of variables (p < 2.2e-16). **(C)** Log2 Odds ratio for the combination of categories shown within each plot, derived from the variables shown on the left. Fisher Exact tests revealed the statistical significance of all enrichments/depletions (p < 1.3e-6). **(D)** The variables shown in (A) were tested as predictors for each region belonging to one of the 3 GATA6 binding groups (Early, Late, Transient). The importance of each feature is shown on the left and the combined predictive success on the right, reaching an Area Under the Curve of 0.81. See Methods for details. 4kb centered on summit

**Supplementary Information, Fig. S 3.**
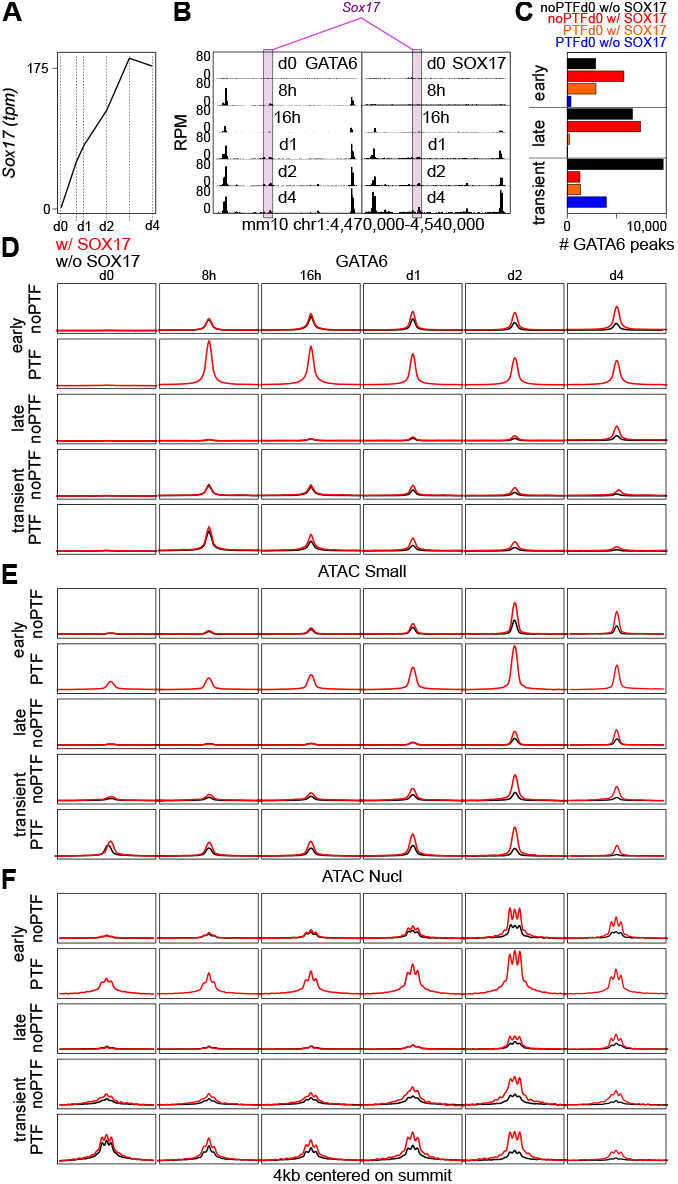
Analysis of SOX17, a target and partner of GATA6. **(A)** RNA-seq expression profile of Sox17 mRNA expressed in Transcripts Per Million (tpm) over GATA6 induction. **(B)** Illustrative co-binding of GATA6 and SOX17 using the extended *Sox17* locus as an example. **(C)** Distribution of GATA6 peaks across different categories based on the dynamics of GATA6 binding (Early, Late, Transient) and the presence/absence of either pluripotency TFs before differentiation (PTFd0/noPTFd0) or presence (w/) or absence (w/o) of SOX17. **(D-F)** The groups capturing most regions in (C) were used to compare GATA6 binding (D), chromatin accessibility using small ATAC-seq fragments (E) and nucleosomes using nucleosome-sized ATAC-seq fragments (F). The plots show the average reads per million across 4kb centered on GATA6 summits.

**Supplementary Information, Fig. S 4.**
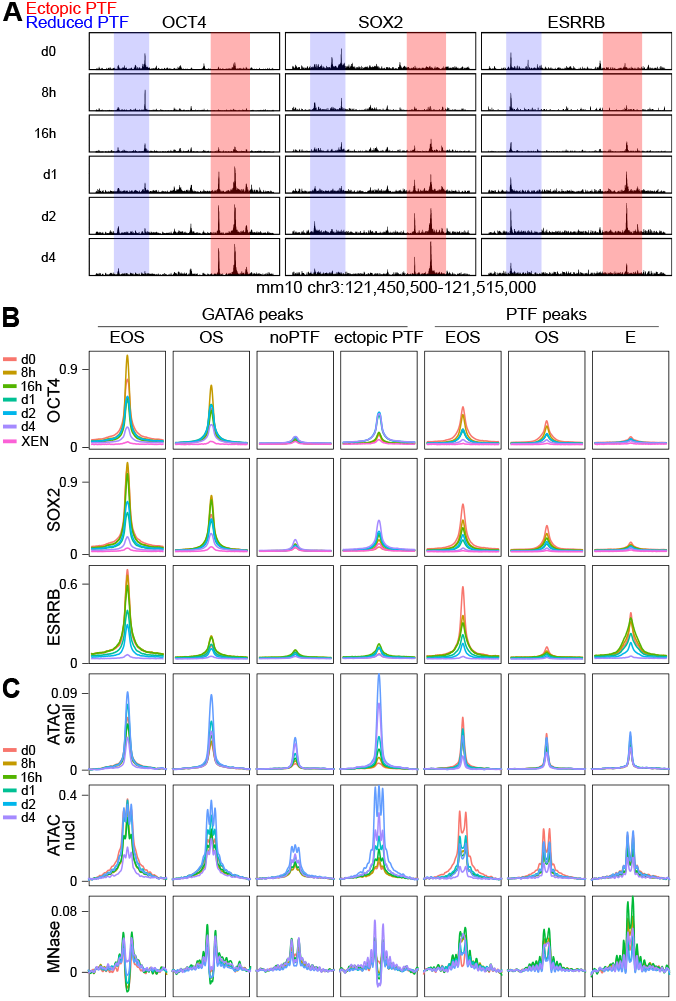
Comparative analysis of GATA6 and pluripotency TF binding regions. **(A)** Example of pluripotency TF binding during GATA6 induction showing regions losing or ectopically gaining their binding. **(B)** Average binding profiles of pluripotency TF at GATA6 and pluripotency TF (PTF) binding peaks throughout all analyzed time-points. Each set of regions was divided according to the binding of either all 3 analyzed pluripotency TFs in undifferentiated cells (EOS for ESRRB, OCT4 and SOX2), only OCT4 and SOX2 (OS), only ESRRB (E), none of the three or when showing ectopic binding during GATA6 induction (ectopic PTF). The regions shown represent 4kb centered on the GATA6 summit for GATA6 peaks or on the summit of chromatin accessibility for PTF peaks. **(C)** Identical analyses for chromatin accessibility measured by small ATAC-seq framents or for nucleosomes measured by nucleosome-sized ATAC-sec or MNase-seq fragments.

**Supplementary Information, Fig. S 5.**
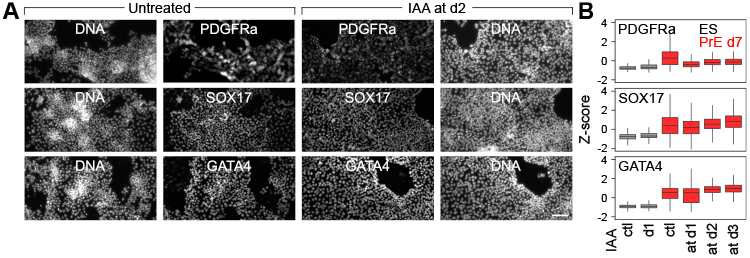
Efficient PrE differentiation upon OCT4 depletion after d1. **(A)** Immuno-staining of PrE markers after 7 days of chemical PrE induction, for untreated (left) and IAA-treated cells from day 2 onwards. The scale bar represents 150 µm. **(B)** Quantification of the immuno-staining of PrE markers in control ES cells (ctl in black), after 1 day of IAA treatment (d1 in black), and in cells differentiated for 7 days (in red) in the absence of IAA (ctl) or having experienced OCT4 depletion from day 1, day 2 or day 3 onwards.

**Supplementary Information, Fig. S 6.**
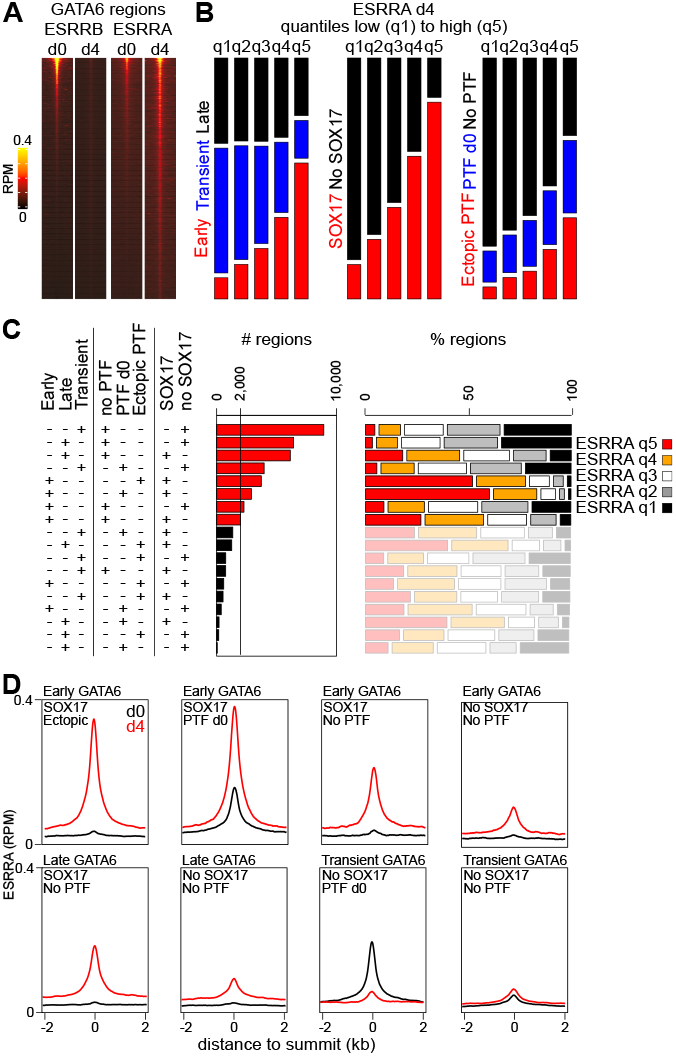
Preferential ESRRA enrichment at functionally relevant GATA6 categories. **(A)** Enrichment levels of ESRRB and ESRRA across all GATA6 binding regions before and 4 days after GATA6 induction, ordered by decreasing ESRRB levels before differentiation. **(B)** Relationships between ESRRA enrichment levels at d4, shown as low (q1) to high (q5) quantiles, and the properties of GATA6 binding regions described before and shown vertically. All 3 combinations of variables were found to be strongly associated (Chi-square test p < 2.2e-16), as were the enrichment of Early or SOX17 or Ectopic PTF at the regions displaying higher ESRRA levels (q5), as assessed with Fisher Exact tests (p < 2.2e-16). **(C)** Repartition of all GATA6 binding regions across all previously studied variables, as shown in the left (18 different combinations). The number of regions within each category is shown on the barplot, as is their composition regarding ESRRA quantiles. All combinations with more than 2000 regions, summing up to 85% of all GATA6 binding regions, were selected for further analyses. **(D)** Average ESRRB binding profile in reads per millions (RPM) before (black) and 4 days after GATA6 induction (red), at the 8 most prominent GATA6 binding regions defined in (C).

**Supplementary Information, Fig. S 7.**
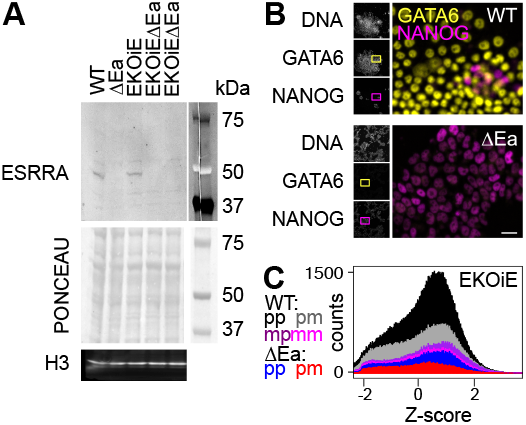
ESRRA and ESRRB requirements for PrE differentiation. **(A)** Western-blot illustrating the loss of ESRRA in ΔEa and in two clones of EKOiEΔEa cells. **(B)** Representative immuno-fluorescence of NANOG and GATA6 after 7 days of PrE chemical induction of wild-type and ΔEa cells. The scale bar represents 20 µm. **(C)** Quantification of the number of EKOiE and EKOiEΔEa cells (Y-axis) expressing different levels of GATA6 (X-axis) after 7 days of PrE chemical induction. ESRRB was depleted by DOX withdrawal either never (pp), concomitantly with the chemical PrE induction (pm), only during 48h before starting the chemical PrE induction (mp), or both 48h before and during the whole PrE chemical induction (mm).

## Notes

### Competing Interest Statement

The authors have declared no competing interest.

