## Supplementary material for "Molecular and epistatic interactions between pioneer transcription factors shape nucleosome dynamics and cell differentiation": Methods

**Materials and methods**

**R.X. Coux et al.**

### a/ Cell Culture

*a1/ General cell culture conditions. Gata6*-inducible ES cells were maintained in ES medium: DMEM (Life Tech 31966047) supplemented with 15% FBS (Life Tech 10270106), 10 ng/mL LIF (Miltenyi Biotech 130-099-895) and 100 µM 2-mercaptoethanol (Gibco #31350-010)) on mitomycin C-treated (Sigma M4287) primary mouse embryonic fibroblasts (feeders). Wild-type E14Tg2a and its derivatives (ΔEa, EKOiE[1] and EKOiEΔEa) were grown in ES medium (10% FBS) on 0.1% gelatin without feeders. OCT4-AID iPS cells[2] were grown in serum-free 2i-containing medium: 1µM PD0325901 and 3 µM CHIR99021 (Axon1408 & Axon1386 respectively), 0.5X DMEM/F12 (Gibco, Cat# 31331093), 0.5X Neurobasal (Gibco, Cat# 21103049), 1X N2 supplement 100X (Gibco, Cat# 17502048), 1X B27 supplement 50X (Gibco, Cat# 17504044), 10µg/ml Insulin (Sigma, Cat# I1882-100MG), 2 mM L-Glutamine (Invitrogen, Cat# 91139), 37.5 µg/ml BSA (Sigma, Cat# A3311-10G), 100 µM 2-mercaptoethanol (Gibco, Cat# 31350-010), 10 ng/ml recombinant LIF (Miltenyi Biotech, Cat# 130-099-895). XEN cells were cultured in DMEM-15% FBS on feeders.

*a2/ Derivation of Gata6-inducible ES cells.* $2\times10^6$ KH2 ES cells[3] were lipofected with 5 µg of PBS31 vector containing the GATA6-3×HA ORF (Ensembl *Gata6-201*, ordered from Genescript) and 2,5 µg of pCAGGS-FLPe (Gene Bridges). The day after, cells were trypsinized and $3{,}5\times10^6$ were seeded on hygromycin resistant feeders with 200 µg/mL hygromycin B. Selection was maintained for 10 days, individual antibiotic-resistant clones were picked, genotyped by PCR and tested for GATA6 induction using 1 µg/mL doxycycline. Two independent clones were selected for further analyses (G6.1 and G6.3).

*a3/ Derivation of ΔEa cells.* E14Tg2a or EKOiE ES cells were nucleofected using an Amaxa Nucleofector II with a Mouse ES Cell Kit (LONZA, Cat# VPH-1001; program A23) with 3 µg of an equimolar pool of two pU6_CBh-Cas9-T2A-mCherry plasmids (Addgene #64324) driving expression of the gRNAs 5'- TCTACCCAAACGCCTCTGCC -3' or 5'- GGGTAGGGGCTCACCCTGGA -3' to induce double strand breaks into the 2nd exon of *Esrra* (Ensembl *Esrra-206*). Two days later single mCherry positive cells were FACS sorted into individual wells of a 96-well plate and expanded. Correctly targeted cells were identified by PCR on genomic DNA and sequencing. Clones bearing deletions on both alleles were selected for further experiments.

*a4/ Primitive endoderm differentiation. Gata6*-inducible cells were seeded at 100,000 cells/cm$^2$ on feeders and differentiation induced by addition of 1 µg/mL doxycycline in ES cell medium, changed daily. Chemical differentiation was induced as previously described[4,5]. Cells were seeded at 37,000 cells/cm$^2$ (ΔEa E14Tg2A and EKOiE cells) or 74,000 cells/cm$^2$ (OCT4-AID iPS cells) onto poly-L-ornithine/laminin coated wells of Ibidi µ-slides (#80446) in endoderm medium: RPMI 1640 Medium, GlutaMAX™ Supplement (Gibco #61870036), 2% B-27 minus insulin (Gibco #15285074) and 100 µM 2-mercaptoethanol (Gibco #31350-010). The day after, the medium was supplemented with 100 ng/ml activin A (R&D # 338-AC-010), 3 µM CHIR99021 (Axon #1386) and 10 ng/ml LIF. The medium was changed every day.

**c/ Imaging.**

*c1/ Bright field microscopy.* Cell culture pictures were taken on a Nikon Eclipse Ti-S inverted microscope equipped with: CFI S Plan Fluor ELWD ×20 objective; 89 North PhotoFluor LM-75; Hamamatsu ORCA-Flash 4.0LT camera; NIS Elements 4.3 software.

*c2/ Immunofluorescence of Gata6-inducible ES cells.* Cells grown on Ibidi µ-slides (#80446) were fixed in PBS-4% PFA (Electron Microscopy Science #15700) for 20 min. Fixed cells were washed 3 times in PBS1X, permeabilized in PBS-0.1% Triton-X100 (Sigma #T8787) for 10 min and blocked for 20 min in PBST – 5% BSA (Sigma #A331). Primary antibody incubation was performed overnight at 4˚C with the following antibodies diluted at 1:1000: goat polyclonal anti-GATA6 (R&D Systems AF1700), mouse anti-OCT4 (clone C10, Santa-Cruz #5279), rabbit polyclonal anti-SOX2 (Active Motif #39823), mouse monoclonal anti-ESRRB (Perseus Proteomics, H6-705-00), goat polyclonal anti-SOX17 (R&D Systems AF1924) and rat monoclonal anti-PDGFRA (APA5, Thermo Fisher #14-1401-82). Secondary antibody incubation was carried out at 1:1000 dilution for 2 hours at RT, with appropriate Alexa Fluor®-conjugated Donkey IgG antibodies (Jackson Immunoresearch). Cells were imaged on a Zeiss LSM800 confocal microscope.

*c3/ Immunofluorescence upon PrE chemical conversion.* Cells differentiated for 7 days were processed into their respective well, washed with PBS1X, fixed with freshly prepared PFA 4% for 10 min at room temperature, washed 3 times in PBS1X, and used for immunostaining. Immunostaining was performed after permeabilization with cold PBS1X/0.5 % v/v Triton X-100 for 5 min, two washes with cold PBS1X, and 30 min incubation on ice with PBS1X/1% Donkey Serum (Sigma, Cat# D9663).

Cells were incubated overnight at 4°C with primary antibodies diluted 1:200 in PBS1X/1% Donkey Serum: goat polyclonal anti-GATA6 (R&D Systems AF1700), rat monoclonal anti-PDGFRA (APA5, Thermo Fisher #14-1401-82), Rabbit Monoclonal anti-SOX17 (Abcam #ab224637), goat polyclonal anti-OCT4 (R&D systems #AF1759), goat polyclonal anti GATA4 (Santa Cruz #sc1237), goat polyclonal anti-OCT4 (Santa Cruz N19 #sc8628), rat monoclonal anti-NANOG (eBioscience #eBIOMLC-51) or rabbit polyclonal anti-NANOG (Cell Signaling Technology #CS8822S). After three washes in cold PBS1X, cells were incubated with secondary antibodies diluted in PBS1X/1% Donkey Serum, washed three times in PBS1X and nuclei counterstained with DAPI (Vectorlabs, Cat#H-1200). Imaging was performed with an inverted Nikon Eclipse X microscope equipped with: X20/0.45 (WD 8.2-6.9) objective; LUMENCOR excitation diodes; Hamamatsu ORCA-Flash 4.0LT camera; NIS Elements 4.3 software. Cell Profiler[6] was used for quantifications.

**d/ Gene expression analyses.**

*d1/ RNA-seq. Gata6*-inducible ES cells were trypsinised at each indicated time-point and feeders adsorbed on non-gelatinized plastic dishes for 20 min. Cells were washed once in PBS1X and lysed in TRIzol. Total RNAs were extracted using chloroform, repurified by acidic pheno-chloroform extraction and precipitated. Stranded, directional polyA selected RNAseq libraries were constructed and sequenced (paired-end 150bp reads) on a NovaSeq6000 system (Illumina) by Novogene.

*d2/ Alignments and quantification.* Stranded paired end reads were aligned to the mouse genome (mm10) using STAR[7], quantified by RSEM[8] with options “ --calc-pme --calc-ci --estimate-rspd --forward-prob 0.0 --paired-end”, and counts converted into Transcripts Per Million (TPM, Table S1) after omitting outliers identified by Principal Component Analysis (> 30 standard deviations from the mean for PC1).

*d3/ Differentially expressed genes (DEGs).* RSEM estimated read counts per sample were rounded and used in DESeq2[9] without independent filtering. All genes with FDR < 0.05 and absolute log2 Fold-Change > 1 in at least one time-point compared to untreated cells (d0) were considered as DEGs. DEGs complying to these filters at 16h or 24h onwards were considered Early_UP/DOWN; at d2 or d3 onwards, Mid_UP/DOWN; at d4 and d7 or only at d7, Late_UP/DOWN. The remaining DEGs displaying more complex behaviors regarding FDR/FC filtering (e.g. FDR < 0.05 at non-consecutive time-points) were clustered based on their expression trends identified using K-means (R function kmeans() with options k=6, nstart=50, iter.max=50). The average profile of each cluster was visualized and the genes attributed to the closest Early/Mid/Late_UP/DOWN group.

d4/ *Developmental annotation.* Correlations to developmental gene expression were made using a published study of mouse embryos around gastrulation[10], with the processed data downloaded from ftp://ftp.ebi.ac.uk/pub/databases/scnmt_gastrulation.

*d5/ Correlations to TF binding sites.* Enrichments of each group of DEGs in proximity to different categories of TF binding sites were calculated using right tail p-values of Fisher Exact tests for the genes of each DEG cluster present within x bp of each TF binding site under consideration, for x in [1:1e+9] bp, confronted to background associations obtained from all considered genes.

**e/ Chromatin assays.**

*e1/ Chromatin Immunoprecipitation and sequencing (ChIP-seq).* Cells were trypsinized, feeders adsorbed, and crosslinked at ~$5\times10^6$ cells/mL in PBS1X supplemented with 2 mM DSG (Sigma-Aldrich, 80424) for 50 min followed by 10 min in PBS1X 1%PFA (Electron Microscopy Science #15700). Crosslinking was stopped with 0.125 mM glycine and cells were pelleted and washed once with cold PBS1X. Crosslinked cells were either used directly for chromatin preparation or frozen in liquid nitrogen and stored at -80℃ for later processing. Fixed cells were resuspended in 2 ml of swelling buffer (25 mM Hepes pH 7.95, 10 mM KCl, 10 mM EDTA, 0.5% IGEPAL and 1× Roche protease cocktail inhibitor) and incubated on ice for 20 min. The suspension was passed > 30 times in a douncer. Nuclei were centrifuged (1500 RPM, 15 min at 4℃) washed and resuspended in TSE150 (0.1% SDS, 1% Triton X100, 2 mM EDTA, 20 mM Tris-HCl pH8, 150 mM NaCl, 1× PIC) at approximately $3\times10^4$ cells/mL. Samples were sonicated in 1.5 ml tubes (Diagenode) using a Bioruptor Pico (Diagenode) for 8 cycles (30s ON / 30s OFF) except for ESRRA ChIP-seq for which a Covaris M220 was used (Setpoint @6°C and 10 cycles of 60 sec duration, peak power of 67W, duty factor of 15% and cycles/burst of 500). After centrifuging at max speed for 15 min, an aliquot was used to check the sonication efficiency and the chromatin stored at -80°C upon further use. Immunoprecipitation was performed with 12,5 µg of chromatin per immunoprecipitation except of ESRRA (20µg), after pre-clearing in TSE150 containing 50 µl of protein G Sepharose beads (Sigma-Aldrich #P3296) 50% slurry blocked with 0,5 mg/mL BSA and 1 mg/mL yeast tRNA (Roche #10109495001) on a rotating wheel for 3 hours at 4˚C. Pre-cleared chromatin was incubated at 4℃ overnight rotating on-wheel with the following antibodies: anti-GATA6 (Cell Signaling technology #5851); anti-OCT4 (abcam ab19857); anti-SOX2 (Active Motif 39823); anti-ESRRB (Perseus

Proteomics, H6-705-00); anti-SOX17 (R&D Systems AF1924), anti-ESRRA (Novus Biologicals NBP147254 and Abcam Ab239879). A 20µL aliquot was reserved for input and 50 µL of blocked protein G beads were added for 1h30 at 4˚C on-wheel. Beads were pelleted and washed on-wheel with 1mL cold buffer in the following order: 3 washes in TSE150, 1 wash in TSE500 (TSE150 but with 500mM NaCl), 1 wash in washing buffer (10 mM Tris-HCl pH8, 0.25M LiCl, 0.5% NP-40, 0.5% Na-deoxycholate, 1 mM EDTA) and 2 washes in TE. ChIP was eluted in 100 mL elution buffer (1% SDS, 10 mM EDTA, 50 mM Tris-HCl pH 8) and beads rinsed in 150mL TE-1% SDS. Eluate and supernatant were pooled. Input and precipitated chromatin was reversed crosslinked overnight (65˚C in TE – 1% SDS with 100 µg stabilized Proteinase K; Eurobio #GEXPRK01), phenolchloroform-extracted and ethanol-precipitated. Libraries were constructed using the NEBNext Ultra II DNA library prep kit (NEB #E7645) using 0.6 µM annealed adapters for ligation. Custom designed Y/forked adapters based on Illumina Truseq indexes with 6-8nt unique molecular identifiers (UMI) 3' of the index sequence were used[11]. Libraries were amplified by 8 to 12 PCR cycles, purified using SPRI beads, quantified with a Qubit 3 (Invitrogen) and fragment size distribution checked on an Agilent 2200 Tapestation. Pooled libraries were sequenced (single end 75bp reads) on a Nextseq2000 sequencing system (Illumina), except ESRRA which was sequenced in PE-mode.

*e2/ Assay for Transposase-Accessible Chromatin using sequencing (ATAC-seq).* 100,000 cells were washed in PBS1X, pelleted and resuspended on ice in 50 µL transposition reaction mix (25 µL 2X TD Buffer, 2.5 µL Tn5 Transposase, 22.5 µL nuclease-free H2O, Illumina Nextera DNA library prep kit). The transposition reaction was incubated at 37˚C for exactly 50 min and stopped by addition of 250 mL of binding buffer (Qiagen MinElute cleanup kit). Transposed DNA was purified with MinElute cleanup kit (Qiagen #28204) and resuspended in 10 µL of H2O. Libraries were prepared by adding 2.5 µL 25 µM primer Ad1.noMX, 2.5 µl 25 µM Ad2 indexing primer[12], 9 µl Nuclease Free Water, 1 µl of a 1:8 dilution of Quant-iT Picogreen dye (Invitrogen, # P11496) and 25 µl of KAPA HiFi HotStart 2× Master Mix (Kapa Bioscience #KK2502) to the transposed DNA. Amplification was carried out in a LightCycler II qPCR machine (Roche) and fluorescence monitored to stop amplification during the exponential phase. Libraries were purified using SPRI beads, quantified using a Qubit 3 (Invitrogen) and fragment size distribution checked on an Agilent 2200 Tapestation. Pooled libraries were sequenced (PE150 reads) on a NovaSeq6000 (Illumina) by Novogene.

*e3/ Micrococcal Nuclease sequencing (MNase-seq).* Cells were fixed in warm ES medium with 1% PFA (Electron Microscopy Science #15700) for 10 min at room temperature and quenched with 0.125M glycine. Cells were pelleted and washed in ice-cold PBS1X. $1.25 \times 10^6$ fixed cells were resuspended in 500µL of MNase buffer (50 mM Tris-HCl pH8, 1 mM CaCl2, 0.2% Triton X-100, 1× Roche Protease cocktail inhibitor) and prewarmed for 10 min at 37°C. 16 U of MNase (NEB Cat# M0247) was added to the reaction and digestion was carried out for exactly 10 min at 37°C with low mixing (300 rpm) on a ThermoMixer (Eppendorf #5384000020). The reaction was stopped on ice by addition of 500 µl of 2× STOP buffer (2% Triton X-100, 0.2% SDS, 300 mM NaCl, 10 mM EDTA). Tubes were placed overnight on a rotating wheel at 4°C to allow diffusion of the digested fragments. The suspension was spun down and the supernatant frozen at -80°C. 25 µL of digested chromatin was reversed-crosslinked overnight (see ChIP-seq), phenol-chloroform-extracted and ethanol-precipitated. Samples were resuspended in TE, incubated with 2 µL of RNAse (Roche #11119915001) for 30 min at 37℃ and purified with SPRI beads. DNA concentration was measured using a Qubit3 (Invitrogen) and size-checked on an Agilent 2200 Tapestation. Libraries were prepared from 20 ng purified DNA: samples were end-repaired, A-tailed for and ligated to 1.25µL of 0.2µM annealed custom adapters as previously described[11]. Libraries were amplified by qPCR and amplification stopped during the exponential phase. Library concentration was measured using a Qubit3 (Invitrogen) and fragment distribution checked on an Agilent 2200 Tapestation. Pooled libraries were sequenced (PE150 reads) by Novogene on a NovaSeq6000 (Illumina).

**f/ Bioinfomatic analyses of chromatin-based datasets.**

*f1/ Reads processing.* Libraries were demultiplexed and fastq files generated (including for index reads for ChIP and MNase-seq libraries) using bcl2fastq conversion software (Illumina). Adapters were trimmed using cutadapt[13] Trimmed reads were aligned with bowtie 2[14] to the mm10 genome using the parameters --local --very-sensitive-local --soft-clipped-unmapped-tlen. Optical duplicates were removed using SAMtools[15] and bams filtered for MAPQ > 30. Reads of bowtie2-aligned SAM files were grouped by names were using SAMtools collate. Mate coordinates and insert size fields of PE reads were filled using SAMtools fixmate. UMIs were added to the SAM RX tag using AnnotateBamWithUmis (https://fulcrumgenomics.github.io/fgbio/). Duplicate reads were marked with SAMtools markdup and filtered. For ATAC-seq, reads were shifted inwards by 4 bp to plus-stranded insertions and −5 bp to minus-stranded insertions, as recommended[12].

*f2/ Peak-calling.* ChIP-seq peaks were called against inputs using MACS2[16] with a q.value cutoff of 0.01 except for GATA6 ($10^{-4}$) and ESRRB (0.05). For each factor and time-point, the bams of all replicates were merged before peak calling and the peaks retained if they were also called in all individual replicates.

*f3/ Peak quantifications.* Quantifications were performed with the bamsignals R package (https://github.com/lamortenera/bamsignals) with systematic correction to the library sizes and counting the number of reads either falling into peak coordinates (Table S2) or covering each base of a 4-kb window centered on the peak summit, defined as the max GATA6 signal for GATA6 peaks or the max ATAC-seq signal for pluripotency TF peaks not bound by GATA6. To identify GATA6 summits the reads were extended as estimated by MACS. For ATAC-seq, PE fragments < 100bp were used to measure accessibility and identify the base with max signal. Quantifications of histone modifications to characterize GATA6 peaks were performed using ES cells bam files downloaded from Encode.

*f4/ DNA motifs.* Motif occurrences and statistics were obtained by scanning GATA6 peaks using FIMO[17] for the following motifs obtained from JASPAR database: MA1104.1 (GATA6), MA0142.1 (OCT4::SOX2), MA0078.1 (SOX17), MA0141.3(ESRRB) and MA0592.1 (ESRRA).

*f5/ Analysis of nucleosomes.* Nucleosomes were mapped either using ATAC-seq or MNase-seq datasets. For ATAC-seq, only fragments of 140:250 bp were considered; for MNase-seq, only fragments of 130:160 bp were considered. The coverage of the signal at specific regions was calculated using fragment midpoints exclusively, normalized to the library depth. The average profiles were smoothed using the loess() function in R, with a span of 0.05 for ATAC-seq and 0.07 for MNase-seq. For MNase-seq data the profiles were internally normalized to the flanking sides (the average of the 1:500,3501:4000 intervals was substracted to the signal). Spectral densities were calculated in R using the spectrum() function and a period of 180 bp, which gave maximal signal.

*f6/ Categorical variables and statistics.* To generate the groups of TF binding regions we used k-means clustering implemented as described before: to split the regions as Early, Transient or Late GATA6 binding sites, we used quantitative data of every time-point of GATA6 enrichment normalized to the corresponding inputs and log2 transformed. For SOX17 (bound by or not) and for pluripotency TFs (bound at d0, ectopically or not at all) we used binary peak calling data. To characterize the regions according to histone modification signatures we used library size normalized, region width normalized and log2 transformed counts, which were subsequently Z-scored for each individual histone mark to scale absolute levels between different marks. The relationships between variables were assessed with the mosaicplot() function in R followed by two-sided Chi-square tests and Fisher Exact tests to compute the odds ratios and associated p-values of specific depletions/enrichments. To test the prediction of GATA6 groups based on different variables we used the Python version of LightGBM[18] for multi-class classification (with max_depth = 10, num_leaves = 180, learning rate = 0.001, lambda_l1: 9.18e-05). Hyperparameters were tuned using a bayesian hyperparameter tuning framework optuna with 5 fold cross-validation (StratifiedKFold(n_splits=5, shuffle=True)). Number of estimators was corrected according to the early stopping option with n=100 and evaluation metric eval_metric = 'multi_logloss'. ROC curve was calculated for the multiclass classification by binarizing the output and implementing macro-averaged one-vs-rest strategy. Feature importance was estimated using pre-build lightgbm attribute feature_importance_ with default parameter importance_type = ‘split’.
